# The relationship between cortisol, grid-like representations and path integration

**DOI:** 10.1101/2025.03.10.642344

**Authors:** Osman Akan, Varnan Chandreswaran, Anne Bierbrauer, Henry Soldan, Nikolai Axmacher, Oliver T. Wolf, Christian J. Merz

**Affiliations:** Department of Cognitive Psychology, Institute of Cognitive Neuroscience, Faculty of Psychology, Ruhr University Bochum, Bochum, Germany; Department of Neuropsychology, Institute of Cognitive Neuroscience, Faculty of Psychology, Ruhr University Bochum, Bochum, Germany; Institute of Systems Neuroscience, Medical Center Hamburg-Eppendorf, Hamburg, Germany

## Abstract

Acute stress triggers the release of cortisol, which broadly affects cognitive processes. Path integration, a specific navigational process, relies heavily on grid cells in the entorhinal cortex (EC). The EC contains glucocorticoid receptors and is therefore likely to be influenced by cortisol, though little is known about this relationship. Given the role of the EC in neurological diseases such as Alzheimer’s Disease, investigating the effects of cortisol on this brain region may offer insights into how stress affects these diseases. In this study, we examined the effects of cortisol on human path integration in thirty-nine healthy participants across two sessions. On each day, they received either 20mg cortisol or a placebo and performed a virtual homing task during functional magnetic resonance imaging (fMRI). Cortisol markedly impaired path integration performance, independent of path distance or the presence of spatial cues. Additionally, cortisol altered navigational strategies, leading participants to navigate further away from landmarks, which was associated with worse performance. FMRI results showed that cortisol increased the activation of right caudate nucleus in the presence of landmarks. Using a representational similarity analysis, we observed grid-like representations in the right posterior-medial EC specifically on day one under placebo, but these were diminished by cortisol. Grid-like representations facilitated performance over short distances but hindered it over longer ones, suggesting that grid cells support PI specifically in case of short trajectories. Overall, the study indicates that cortisol-induced disruption in grid cell function in the EC may underly stress effects on path integration.

## 1 Introduction

Finding ourselves in situations where we lost orientation can easily invoke feelings of discomfort or stress, but we also often experience the inverse relationship of stress causing spatial disorientation, mainly through altering the way we navigate. For example, under time pressure, we tend to use more familiar and less accurate trajectories (1). In general, however, human spatial navigation has not gained much attention in the realm of stress-related consequences, albeit being a type of cognition that is strongly based on hippocampus (HC) and adjacent cortical regions and thus a major target of stress hormones. Investigating the effects of stress on navigation may ultimately deepen our understanding of neurological diseases, particularly since navigational deficits could serve as early biomarkers for Alzheimer’s Disease (2), and stress has been linked to both its pathogenesis and progression (3).

The stress response involves the activation of the rapid sympathetic-adrenal-medullary axis (SAM) and the slow hypothalamic-pituitary-adrenal axis (HPA), generally conceptualized as an adaptive reaction (4). SAM axis activity initiates the release of catecholamines (noradrenaline and adrenaline), while HPA axis activity leads to the release of glucocorticoids (cortisol in humans, corticosterone in most rodents). Cortisol effects are mainly exerted by binding to mineralocorticoid receptors (MRs) and glucocorticoid receptors (GRs), while noradrenaline binds to adrenergic receptors (5, 6). Thereby, acute stress or pharmacological cortisol administration affect several cognitive domains including executive functions, episodic memory, fear conditioning, and decision making (7–10).

During spatial navigation, different strategies can be applied depending on several factors including environmental features (11). One strategy is path integration (PI), a process mainly used in absence of spatial cues and for short trajectories because error accumulation makes it less suited for longer trajectories. PI involves integration of self-referential information to estimate the current position and orientation in relation to an arbitrary reference point (12). On the neuronal sheet, PI has been related to head-direction cells (13) and grid cell firing in the entorhinal cortex (EC; 14–17). Specifically, the characteristic arrangement of grid cell firing fields in a regular hexagonal pattern (18, 19) may provide a general spatial metric of distances (20). In presence of spatial cues, however, additional neural systems tuned to the specific type of cue (e.g., boundary or landmark) are recruited, including HC (21) and posterior cingulate/retrosplenial cortex (PC/RSC; 22). These additional systems may either stabilize grid cell firing (23) or support complementary navigational strategies. Because processing in the striatum, especially the caudate nucleus (CN), is involved in landmark-based navigation (24) and appears to be enhanced under acute stress (25), the presence of spatial cues likely moderates stress hormone effects on either performance or strategy use during navigational processes.

Crucially, the EC is an important mediator of the stress response itself (26–29) and the abundance of GRs in the EC (30) as well as its interdependence with the HC makes it a likely target of cortisol effects. Accordingly, acute stress impairs PI over long distances and in environments with little spatial information (31), and chronic stress is associated with impaired PI under similar conditions (32). These effects might reflect cortisol-induced alterations in grid cell activity in the EC. Furthermore, stress appears to alter navigational patterns, leading participants to opt for routes closer to a landmark (31). This suggests an increased recruitment of landmark-based strategies under stress, possibly due to enhanced striatal processing.

In this study, we aimed to investigate whether cortisol administration mimics the effects of stress on PI and to uncover the underlying neural mechanisms using functional magnetic resonance imaging (fMRI). To assess PI performance, we used an adapted version of the “Apple Game” paradigm implemented by Bierbrauer et al. (22). In different subtasks of this virtual PI task, participants can either rely only on visual flow (Pure PI) or on visual flow and a central landmark (Landmark PI; Fig. 1). To measure proxies of grid cell activity, we analyzed so called “grid-like representations” (GLRs; 33, 22, 34).

**Figure 1.**
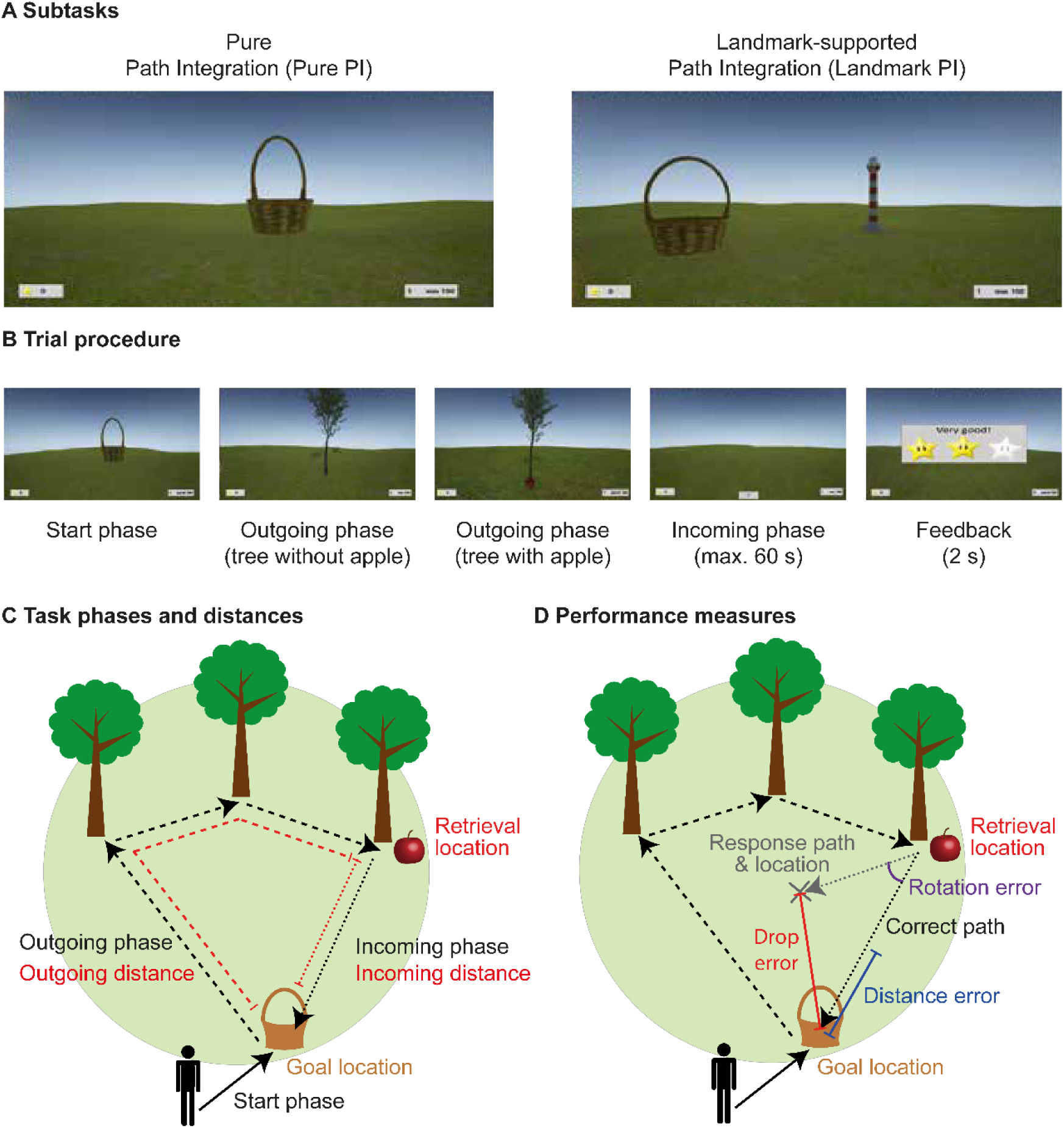
Experimental path integration task. (A) While the Pure PI subtask consisted only of a grassy plain, the Landmark PI subtask additionally contained a central lighthouse serving as spatial cue. (B) Each trial began with the “start phase”, where participants navigated to a basket (goal location), the location of which they should encode. In the following “outgoing phase”, they navigated to a variable number of trees (1–5) until reaching a tree containing an apple (retrieval location). Then, during the “incoming phase”, participants had to find the way back to the goal location before receiving feedback via zero to three stars according to performance based on the drop error (see D). Basket and trees disappeared as soon as they were reached. (C) Outgoing phase (dashed black line) and incoming phase (dotted black line) were quantified according to their spatial distances: outgoing distance corresponded to the cumulated distance from goal to retrieval location (dashed red line), and incoming distance to the Euclidean distance between retrieval and goal location (dotted red line). (D) General PI performance was assessed via the drop error, which corresponded to the distance between response location (marked with an X) and goal location (solid red line). The drop error can further be differentiated into distance error, referring to the difference between retrieval-to-goal distance and retrieval-to-response distance (blue line), and rotation error, depicting the angle between the retrieval-to-goal path and the retrieval-to-response path (purple arc). Figure adapted from Bierbrauer et al. (22).

Our results strongly support the hypothesis that stress-induced deficits of PI are mediated via cortisol, but they do not show that cortisol changes the navigational pattern towards the landmark. The fMRI data demonstrated the importance of PC and CN during Landmark PI and for the CN during Landmark PI under cortisol. Additionally, they showed that the EC is most relevant during Pure PI under placebo. Importantly, our findings suggest that cortisol impairs GLRs, which were particularly beneficial for PI over short distances.

## 2 Results

We analyzed data from 39 men aged 19 – 34 years (24.18 ± 4.55 years; mean ± SD) participating in a two-day within-subject crossover design. Participants received either 20 mg of cortisol or visually identical placebo tablets on day one, and the respective other treatment on day two (one week later), before completing the Apple Game during fMRI recordings on both days. The order of administration was counterbalanced across participants (Fig. 2; STAR Methods). To examine effects of cortisol on PI, we conducted univariate and multivariate fMRI data analyses and built several linear mixed models (see Tab. S1 for an overview). In all mixed models, “subject” was added as random factor, and age, testing day, and sequence (to control for order effects) as covariates.

**Figure 2.**
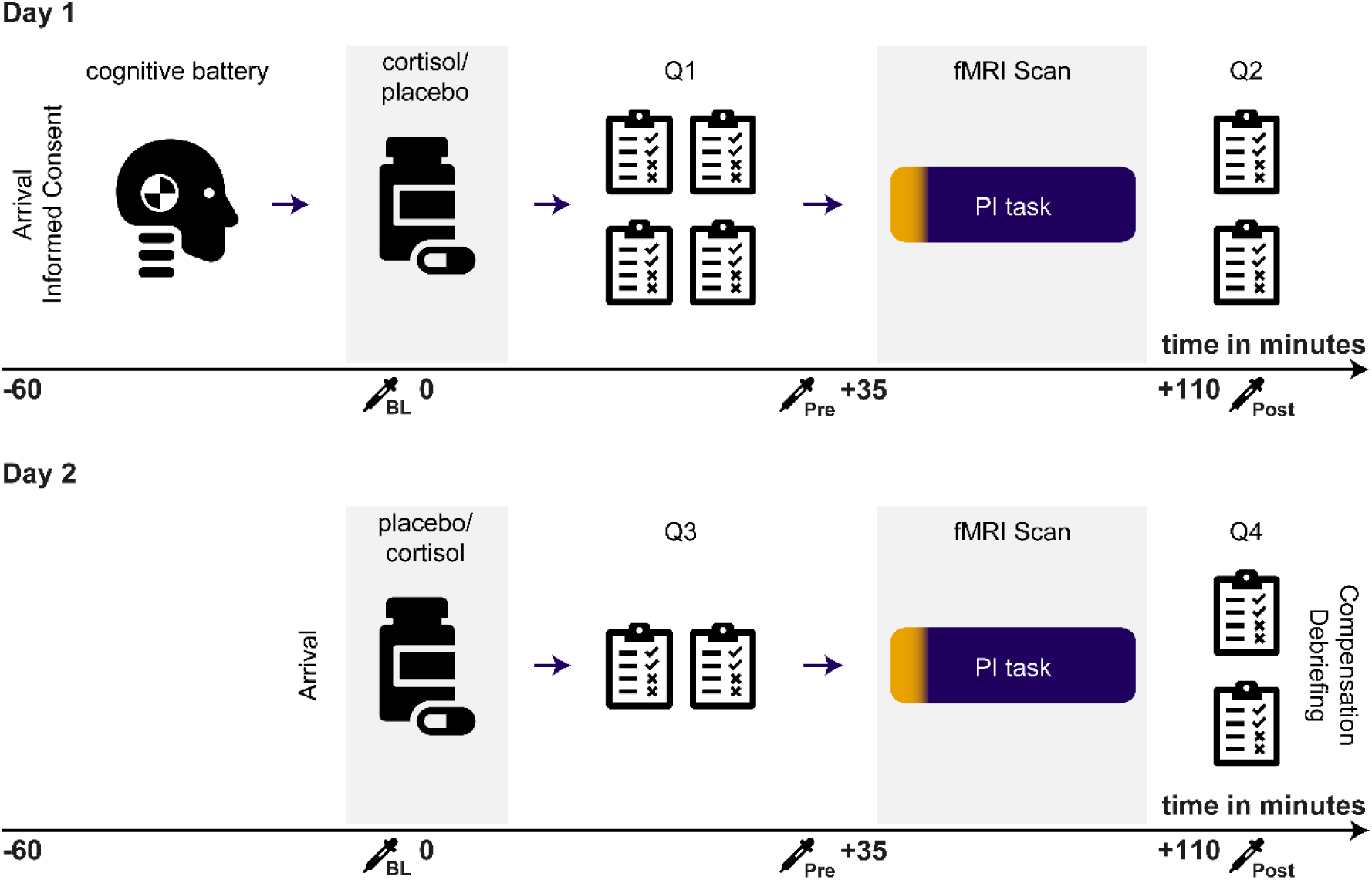
Experimental procedure. All participants were tested in both conditions (cortisol and placebo) on two testing sessions with an inter-session interval of one week. At the beginning of day one, participants underwent a cognitive test battery, which lasted about 40 minutes. Participants then received either 20 mg of cortisol, or a placebo. Afterwards, questionnaires (Q1) assessing demographic, psychological and medical data were filled out. Participants then were prepared for the fMRI scan and the PI task started approximately 40 minutes after cortisol/placebo administration, before answering final questionnaires including a treatment guess and assessment of individual navigational strategies (Q2). One week later, participants were invited for the second testing session. The procedure on day two was similar, except that it did not contain a cognitive test battery, included less questionnaires and involved the respective other pharmacological intervention as compared to day one (crossover design). Numbers on x-axes reflect time-points relative to tablet administration, 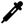 collection of saliva sample, PI: path integration.

### 2.1 Cortisol impairs PI performance irrespective of path distance and availability of spatial cues

The analysis regarding the success of cortisol administration revealed elevated cortisol concentrations under cortisol compared to placebo treatment for the entire PI task (Tab. S2, STAR Methods). In the PI task, we assessed different measures for path distance, PI performance and navigational pattern during the experimental task (STAR Methods). For path distance, we focused on *incoming distance*, which refers to the Euclidean distance between retrieval and goal location (Fig. 1C). For overall PI performance, we assessed the drop error, i.e., the Euclidean distance between response location and goal location (Fig. 1D). Moreover, to investigate the specific role of the landmark under cortisol, we measured the distance between the goal and the spatial cue (goal-to-landmark distance), and the mean Euclidean distance of the moving participant from the landmark across all time points of the incoming phase (movement-to-landmark distance).

To investigate whether cortisol influences PI performance, we built a linear mixed model, in which we investigated effects of subtask (two levels: Pure PI vs. Landmark PI), path distance (i.e., incoming distance), and treatment (two levels: cortisol vs. placebo) on overall PI performance (i.e., drop error) on the level of single trials. This analysis revealed similar findings as in our previous work (31, 32, 22): We observed main effects of subtask (*F*(_1,4945_) = 432.88, *p <* .001, η_p_^2^ = .080), and of incoming distance (*F*(_1,4945_) = 364.99, *p <* .001, η_p_^2^ = .069), indicating higher errors in Pure PI than Landmark PI and for longer incoming distances, respectively. Besides, we found an interaction effect between subtask and incoming distance (*F*(_1,4949_) = 44.99, *p <* .001, η_p_^2^ = .009; Fig. 3A, left), indicating a stronger relationship between incoming distance and drop error in Pure PI as compared to Landmark PI (*t*(_4949_) = 6.71, *p <* .001, *d* = 0.095). Moreover, we found a main effect of age (*F*(_1,36_) = 12.23, *p =* .001, η_p_^2^ = .254; Fig. 3A, right), reflecting higher drop errors in older age. No effects of testing day (*F*(_1,4945_) = 0.39, *p =* .532, η_p_^2^ < .001) or sequence (*F*(_1,36_) = 2.18, *p =* .148, η_p_^2^ = .057) emerged, indicating the absence of an order effect. Importantly, we found a main effect of treatment on drop error (*F*(_1,4945_) = 10.46, *p =* .001, η_p_^2^ = .002; Fig. 3B, left), indicating that cortisol impaired overall PI performance. Cortisol did not interact with other variables, suggesting a general effect independent of spatial cues or path distance.

**Figure 3.**
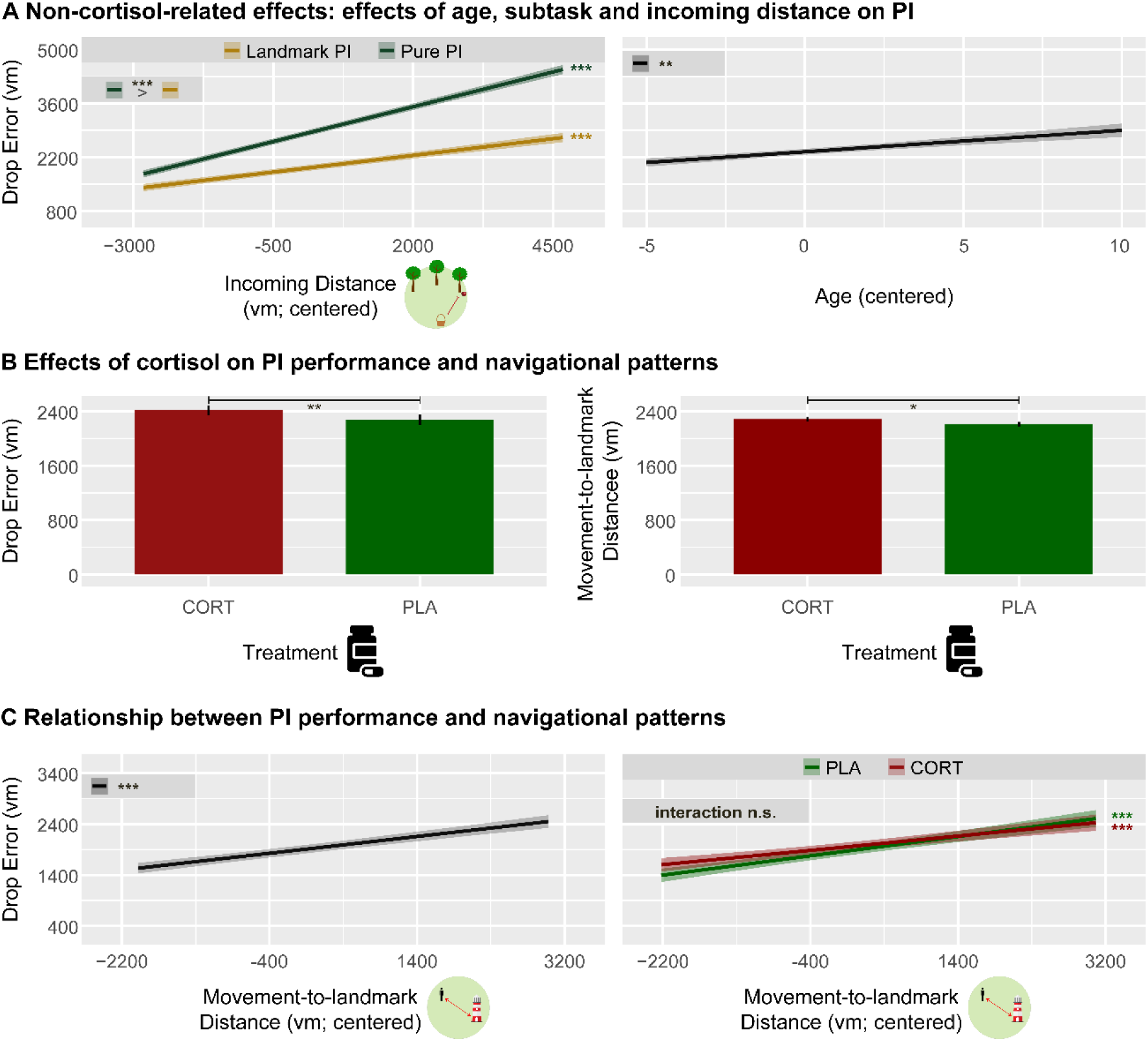
Predictors of PI performance and strategy use. (A) The effect of longer incoming distances leading to higher drop errors was more pronounced when no spatial cues were available (left), and older age led to higher drop errors (right). (B) CORT led to overall higher drop errors (left), and to navigating further away from the landmark in Landmark PI (right). (C) In Landmark PI, the movement distance to the landmark predicted performance, irrespective of treatment. Significant slopes are indicated via asterisks. Error bars and confidence bands represent SEM. Pure PI: pure path integration, Landmark PI: landmark-supported path integration, CORT: cortisol, PLA: placebo, vm: virtual meters, n. s.: not significant, \*\*\**p* < .001, \*\**p* < .01, \**p* < .05.

In a second step, we built linear mixed models only including Landmark PI to examine the navigational pattern in the presence of the landmark under heightened cortisol levels. These models aimed at investigating the role of goal-to-landmark distance and its interaction with cortisol on PI performance, and at investigating whether cortisol affected the employment of navigational strategies (irrespective of performance), respectively. In the first model, we found that a longer distance between goal location and landmark predicted worse PI performance, and cortisol did not moderate this effect (*F*(_1,2453_) = 348.20, *p* < .001, η_p_^2^ = .124). Like in the main models, older age again significantly reduced PI performance (*F*(_1,36_) = 6.30, *p* = .017, η_p_^2^ = .149). In the second model, importantly, we found a main effect of treatment on movement distance to the landmark (*F*(_1,2455_) = 4.57, *p* = .033, η_p_^2^ = .002; Fig. 3B, right), showing that cortisol treatment led participants to navigate further away from the landmark, suggesting that they used landmark information less than under placebo.

The behavioral results so far demonstrated that cortisol impairs PI performance and alters navigational strategy toward navigating further away from landmarks (if available). To analyze whether these alterations were related to each other, we conducted a further analysis examining the effect of movement distance to the landmark on the drop error. Indeed, navigating further away from the landmark was detrimental for performance (*F*(_1,2453_) = 33.46, *p* < .001, η_p_^2^ = .013 ;Fig. 3C, left), and this effect was equally observed under placebo and under cortisol as evidenced by a lack of an interaction effect (*F*(_1,2463_) = 0.72, *p* = .395, η_p_^2^ < .001 ;Fig. 3C, right). The cortisol-induced change in navigational strategy may thus have contributed to the overall detrimental effect of cortisol on PI performance.

### 2.2 Cortisol enhances activation in the right CN in the presence of landmarks

We tested the contrasts “Landmark PI > Pure PI”, “CORT > PLA” and their interaction “(Landmark PI > Pure PI)_CORT_ > (Landmark PI > Pure PI)_PLA_” – as well as the respective reverse contrasts – during the outgoing and incoming phases. We performed an exploratory whole-brain analyses and ROI analyses in left and right HC, left and right CN, bilateral PC, and left and right pmEC for each contrast. For the whole-brain analysis, statistical parametric maps were initially thresholded at a family-wise error (FWE)-corrected α level of *p* < 0.05 across the whole brain and clusters were considered significant at *p* < 0.05, FWE-corrected (extent threshold of five voxels). For ROI analyses, the significance threshold was set to *p* < 0.05 on voxel level, corrected for multiple testing within each ROI using FWE-correction with the small volume correction option of SPM12.

The exploratory whole-brain analyses revealed significantly higher activation in left superior parietal lobule, bilateral precuneus (adjacent to PC) as well as in visual areas including superior and middle occipital gyri during Landmark PI compared to Pure PI, presumably reflecting the higher amount of visual information (see Fig. S2 and Tab. S3). No significant clusters were found on whole brain level for the other contrasts.

The ROI analyses (see Fig. S3 for overview of masks) showed higher activation of PC (*t*(_35_) = 6.33, *p*_FWE_ *<* .001) and right CN (*t*(_35_) = 4.45, *p*_FWE_ *=* .010) in Landmark PI compared to Pure PI (*LPI > PPI*; Fig. 4A), while no significant differences were found for the other ROIs. Furthermore, no ROI showed differential activation in the contrast between cortisol and placebo (*CORT > PLA*). Importantly, however, we observed higher activation in the right pmEC (*t*(_35_) = 2.78, *p*_FWE_ *=* .045), right CN (*t*(_35_) = 4.62, *p*_FWE_ *=* .006), and left CN (*t*_(35)_ = 3.90, *p*_FWE_ *=* .035) in the interaction contrast of Landmark PI vs. Pure PI during cortisol compared to placebo (*[Landmark PI > Pure PI]_CORT_ > [Landmark PI > Pure PI]_PLA_*).

**Figure 4.**
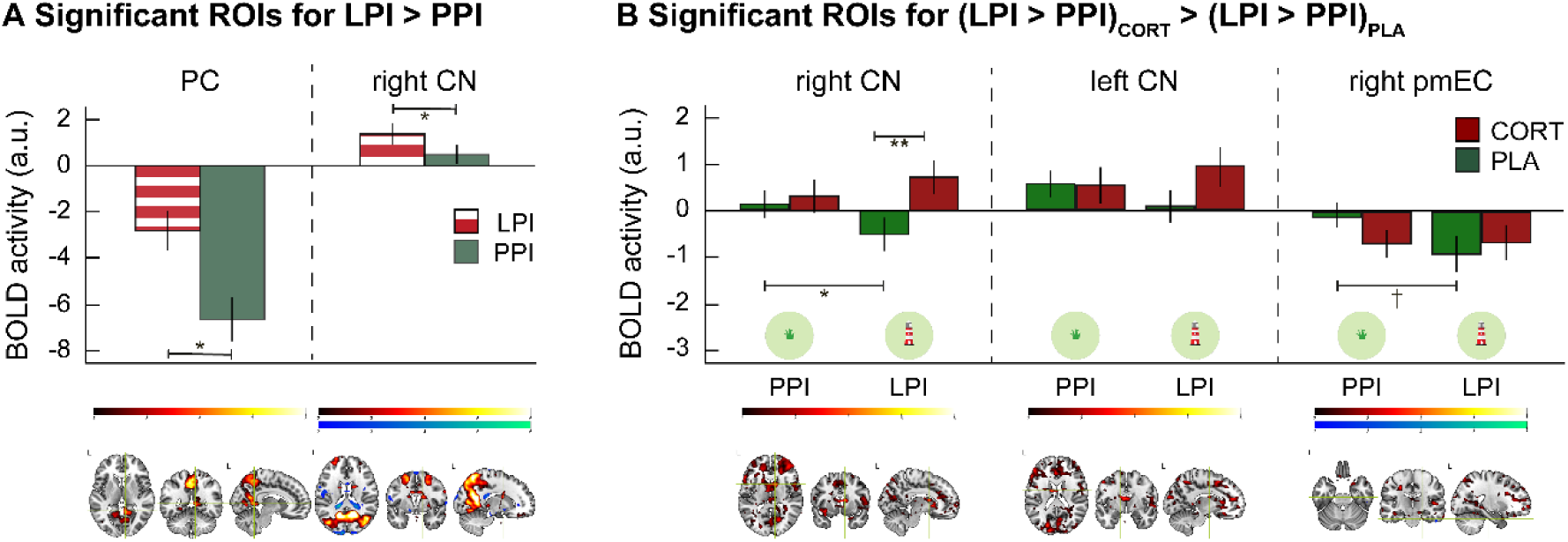
Univariate fMRI results. (A) Significantly higher activation in PC and rNC during LPI as compared to PPI. (B) Significantly higher differentiation between subtasks under different treatments or between treatments in different subtasks in right CN, left CN and right pmEC. Error bars represent SEM. PC: posterior cingulate, CN: caudate nucleus, pmEC: posterior-medial entorhinal cortex, PPI: pure path integration, LPI: landmark-supported path integration, CORT: cortisol, PLA: placebo, n.s.: not significant, ** *p* < .01, * *p* < .05, ^†^ *p* < .10.

Pairwise post-hoc tests showed that these interactions reflected relatively higher activation of the right CN under cortisol compared to placebo during Landmark PI (*t*_(35)_ = 3.43, *p*_Bonferroni_ *=* .006, *d* = 0.550) and during Pure PI compared to Landmark PI under placebo (*t*_(35)_ = 2.82, *p*_Bonferroni_ *=* .032, *d* = 0.291), as well as a trend for a higher activation of the right pmEC during Pure PI than Landmark PI under placebo (*t*_(35)_ = 2.45, *p*_Bonferroni_ *=* .077, *d* = 0.366). No significant pairwise comparisons emerged for left CN (all *t* ≤ 2.29, all *p*_Bonferroni_ ≥ .112, all *d* ≤ 0.360; Fig. 4B). These findings indicate a reduced recruitment of right CN during Landmark PI compared to Pure PI under placebo, but enhanced recruitment during Landmark PI under cortisol compared to placebo, suggesting a relatively increased relevance of this region in the presence of landmarks under pharmacologically induced stress. Conversely, the findings suggest that the presence of landmarks can reduce recruitment of the right pmEC. No other ROI exhibited higher activation for this contrast, neither for any of the reverse contrasts.

### 2.3 Cortisol impairs GLRs, which benefit PI performance specifically over short distances

To assess GLRs, we used a multivariate approach and focused on right pmEC, consistent with previous studies (33, 22). Accordingly, due to their sixfold rotational symmetry, grid cells should show similar activity during movements along directions that differ by 60° (aligned movements) as compared to those differing by 30° (misaligned movements; STAR Methods).

We first aimed to confirm the presence of GLRs by comparing activity during aligned vs. misaligned movements (Fig. 5A) and then tested whether they were affected by cortisol. Indeed, we found higher similarities between aligned movements (mod(α,60°) = 0°) than misaligned movements (mod(α,60°) = 30°) across all days and treatments, demonstrating the presence of GLRs in our study (*t*_(71)_ = 2.92, *p =* .002, *d* = 0.345, Fig. 5B, top-left). Subsequent analyses revealed no main effect of treatment on GLRs (*F*_(1,67)_ = 0.15, *p* = .696, η_p_^2^ = .002), but we observed a significant treatment x day interaction (*F*_(1,67)_ = 6.79, *p* = .011, η_p_^2^ = .092; Fig. 5B, top-right). Due to the crossover design, the interaction cannot easily be interpreted, as it may also include order effects. We thus decided to examine differences between groups on day one (35). When doing this, post-hoc tests revealed a significant effect of treatment (*t*_(67)_ = 2.01, *p =* .040, *d* = 0.700), indicating significantly higher GLRs during PLA than during CORT. Indeed, separate analyses confirmed the presence of GLRs under placebo (*t*_(18)_ = 2.38, *p*_Šídák_ *=* .029, *d* = 0.545, Fig. 5B, bottom-right) but not under cortisol treatment (*t*_(16)_ = 0.20, *p*_Šídák_ *=* .665, *d* = 0.049). Control analyses of GLRs from day one under placebo (Fig. 5B, bottom-right) confirmed that pattern similarity differed between aligned and misaligned movements only for 6-fold rotational symmetry, but not for 4-fold, 5-fold, 7-fold, or 8-fold rotational symmetries (all t ≤ 1.91, all p ≥ .073), and removing movements during same heading directions (±15° from 0°) preserved (by trend) GLRs (*t*_(18)_ = 1.57, *p =* .067, *d* = 0.36).

**Figure 5.**
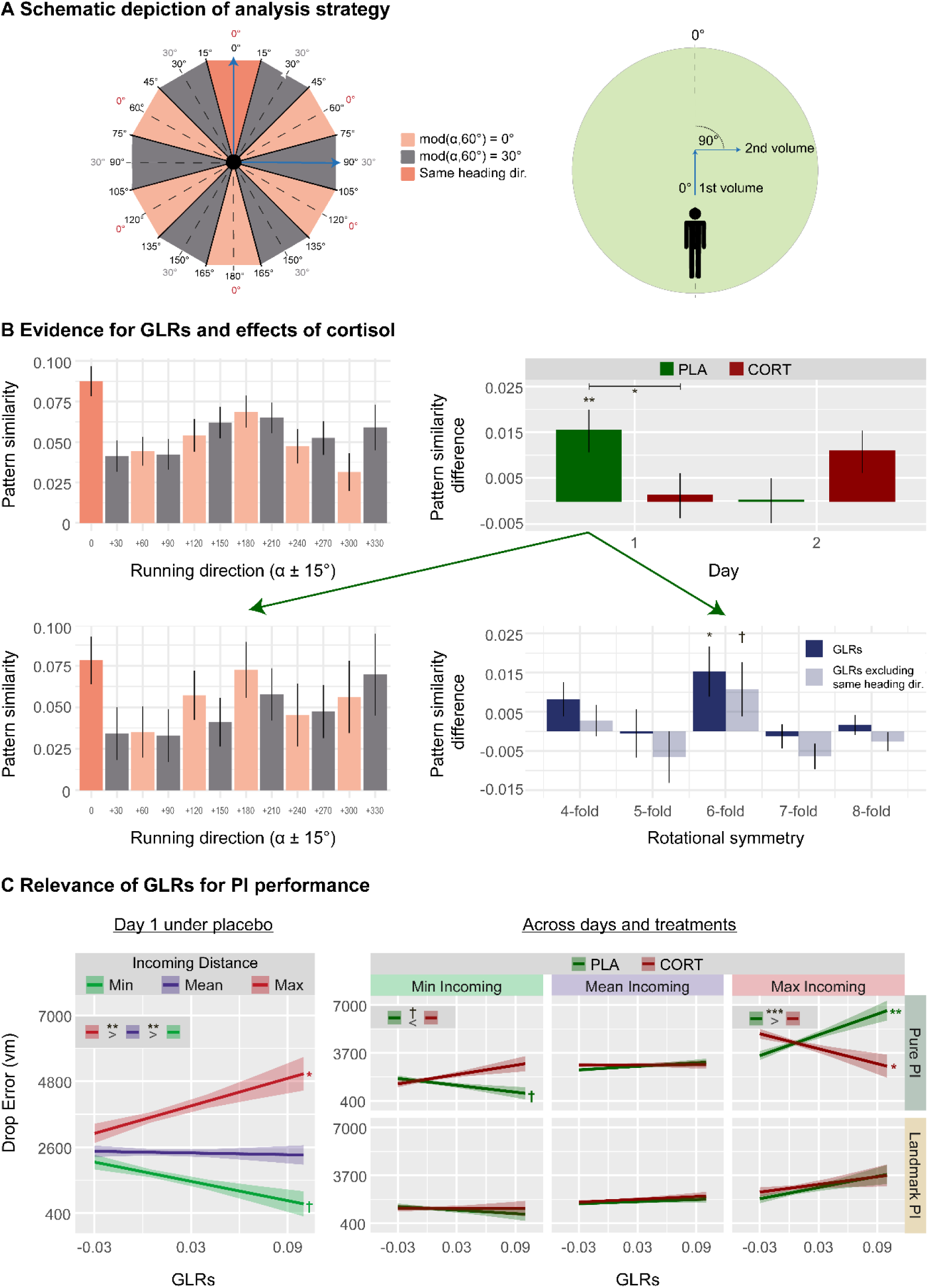
Cortisol effects on GLRs and their behavioral relevance. (A) Inner numbers represent angular differences in 360° space, whereas outer numbers show differences in 60° space (left panel). Higher pattern similarity was expected for angular differences of mod(α,60°) = 0° (rose; α as angular difference between two movement directions) than for angular differences of mod(α,60°) = 30° (gray). The same result was expected when discarding pattern similarities of movements for the same heading direction (dark rose). Two exemplary fMRI volumes representing different movement directions (blue arrows; right panel). In the first volume, the participant navigates at an angle of 0° with respect to the (arbitrary) reference axis, while navigating at an angle of 90° in the second volume. This results in an angular difference of 90° in 360° space, corresponding to 30° in 60° space and thus mod(α,60°) = 30° (compare blue arrows on the left panel). (B) Average pattern similarity over the entire time series of all voxels in the right pmEC for all directions of aligned (rose) and misaligned (grey) fast runs (top-left panel). Differences in pattern similarity depending on treatment and day (top-right panel), and average pattern similarity (bottom-left panel) as well as control analyses (bottom-right panel) only for PLA treatment on day one. (C) Relevance of GLRs for PI considering day 1 under placebo (left) and across days and treatments (right). Significant slopes are indicated via asterisks. Min Incoming, Mean Incoming, and Max Incoming refer to the minimum, mean and maximum incoming distances, respectively, and were chosen as basis for estimating adjusted means of linear trends. Error bars and confidence bands represent SEM. GLR: grid-like representation, pmEC: posterior-medial entorhinal cortex, Pure PI: pure path integration, Landmark PI: landmark-supported path integration, CORT: cortisol, PLA: placebo, vm: virtual meters, n.s.: not significant, ** *p* < .01, * *p* < .05, ^†^ *p* < .10.

Subsequently, we investigated whether GLRs were behaviorally relevant using a linear mixed model incorporating effects of subtask, path distance, and GLRs. Because we only found GLRs on day one under placebo, these analyses focused on this condition. We found similar effects as in the purely behavioral model (main effects of subtask, incoming distance, age, and the subtask x incoming distance interaction) with the addition of an interaction between incoming distance and GLRs (*F*_(1,1193)_ = 10.47, *p* = .001, η_p_^2^ = .009; Fig. 5C, left). Post-hoc analyses revealed that GLRs are (by trend) beneficial for performance (lower drop errors) in case of short incoming distances (*t*_(44)_ = 2.29, *p*_Šídák_ *=* .079), while they were detrimental for long incoming distances (*t*_(133)_ = 2.45, *p*_Šídák_ *=* .046). For the sake of completeness, we repeated this analysis considering data across days and treatments, which yielded in a more complex four-way interaction between GLRs, subtask, incoming distance and treatment (*F*_(1,4560)_ = 5.29, *p* = .021, η_p_^2^ = .001; Fig. 5C, right). This interaction suggested that the effects of GLRs are limited to environments without spatial cues (pure PI). More specifically, during pure PI under placebo, GLRs tended to be beneficial for short distances, but detrimental over long distances, similar to the findings from day 1 under placebo. In contrast, during pure PI under cortisol, GLRs were beneficial over long distances. However, this last analysis incorporating all data should be interpreted with caution, as GLRs were only reliably observed on day 1 under placebo.

### 2.4 Exploratory parametric modulation analysis of PI performance

We further conducted a parametric modulation analysis using a general linear model (GLM), including trial-wise inverted drop error values as regressor for the combined outgoing and incoming phases. We did not distinguish between subtasks or treatments as we were interested in general performance-related neural activity across conditions, encompassing both whole-brain analysis and ROI analyses. The whole-brain analysis yielded a set of clusters showing activation modulated by trial-wise drop error, while the ROI analyses did not reveal any significant modulation of the BOLD signal (all *p*_FDR_ *>* .630; Fig. S4 and Tab. S4).

## 3 Discussion

We investigated whether cortisol administration affects PI performance, navigational patterns in the presence of a landmark, and associated brain activations in healthy young participants. We found cortisol to impair overall PI performance and to promote a change in navigational pattern towards navigating further away from the landmark. Because navigating further away from the landmark was linked to worse performance, this cortisol-induced change in navigational pattern likely contributed to the observed PI deficits. The univariate fMRI data analysis indicated a higher activation of PC and right CN when PI was supported by landmarks, and of right CN in this condition under cortisol. In the absence of landmarks, when participants could solely rely on optic flow, right pmEC showed relatively higher activation under placebo compared to cortisol. The multivariate fMRI data analysis showed the presence of GLRs in the right pmEC under placebo on day one, but not under cortisol, and while they were (on trend) beneficial for PI performance over short distances, they were detrimental over long distances.

Our behavioral results showed that cortisol impairs overall PI performance, an effect mediated by slight impairments of distance and rotation estimations (see Fig. S5). Even though we expected a detrimental cortisol effect, we assumed it to be more pronounced in environments with no spatial cues (Pure PI) and in trials with longer incoming distances, because we had previously found acute stress effects to be moderated by these variables (31). Cortisol is a relevant mediator of stress effects and known to influence cognitive processes (8), but because its increase is only one part of the several biological processes involved in the stress response, effects of cortisol and acute stress are often not identical and sometimes even opposing (36). An important principle of glucocorticoid effects is dose-dependency (37), referring to differential effects on neurons in a concentration-depending manner. Dose-dependency interacts with the local distribution pattern of MRs and GRs, which exhibit differences in their affinity to cortisol (5). While MRs show greater affinity and are highly expressed in limbic structures, GRs possess lower affinity and are more uniformly distributed across the brain, with high abundance in the EC (30). Thus, high cortisol concentrations like in the current study (much higher than average acute stress responses) may affect GRs in EC more strongly and thus impair PI irrespective of moderators like environmental scarcity and path distance. In another previous study, we did not find a main effect of chronically elevated cortisol levels on PI performance, but subtle impairments of distance estimations under conditions of spatial scarcity and long path distance (32). Again, cortisol concentrations were in a low-to-mid range, potentially being insufficient to occupy enough GRs to impair PI performance generally. We therefore propose that cortisol affects PI in a concentration-dependent manner, where lower concentrations lead to subtle effects, detectable in the absence of spatial cues and over long path distances, and where higher concentrations induce severe effects, detectable irrespective of environmental properties or path distance.

The role of landmarks during PI is to stabilize grid cell firing (23, 38), thereby improving performance in general, an effect that is in accordance with our results. When examining the navigational pattern during Landmark PI irrespective of performance, however, we surprisingly found that cortisol led to navigating further away from the landmark. Notably, this effect may partly explain the cortisol-induced impairment in overall PI performance, because increased distances to the landmark generally predicted higher drop errors. Conversely, in a previous study, we had shown that acute stress (on trend) changes the navigational pattern towards the landmark, and we argued that this change might reflect stress-induced enhanced use of landmark information, stronger recruitment of the PC/RSC, or a shift towards striatal processing after acute stress (31). While the fMRI data of our current study do not support the idea of higher recruitment of PC/RSC, we indeed found higher activation in right CN under cortisol during Landmark PI compared to Pure PI, supporting the claim of increased striatal processing. Mechanistically, acute stress enhances the use of landmark information in the striatum via cortisol binding on MRs (39), especially in participants showing high cortisol responses. Thus, we would here expect converging effects of acute stress and cortisol, even though dose-dependency might again play a relevant role. With high cortisol concentrations, like in our current study, not only MRs are activated in the striatum, but also GRs, changing the relative activation of MRs and GRs and thereby altering neuronal processing, an idea referred to as *MR/GR ratio hypothesis* (40). However, while the potential concomitant activation of both receptor types in this study might have contributed to the inconsistency in findings, our task is not specifically designed for investigating landmark-based strategies and their neuronal correlates, making it less comparable to studies targeting this phenomenon explicitly. Further research is needed to clarify the role of cortisol (concentration) on landmark use, both in general and specifically during PI.

The exploratory whole-brain analyses demonstrated higher activation in the left superior parietal lobule, bilateral precuneus and in visual areas in the presence of landmarks. The precuneus provides spatial information to HC and EC (41) and its higher activation along with that of visual areas during Landmark PI presumably reflects the higher availability of visual information. Additionally, the parametric modulation analysis of the drop error revealed that these brain regions were also related to PI performance (among others). The ROI analyses did not reveal any region that is modulated by PI performance, but showed less deactivation of the PC and higher activation of the right CN during Landmark PI, supporting the hypothesis of their crucial involvement in landmark processing (42, 38, 43, 22). Furthermore, we did not observe any ROI that shows generally more activation under cortisol than placebo, but when incorporating subtask, we found effects in right CN and right pmEC. Right CN showed higher activation under cortisol during Landmark PI, suggesting that striatal processing might not be generally enhanced under stress, but specifically when usage of stimulus-response type of strategies (landmark processing in this case) is possible. Under placebo, the right pmEC showed less deactivation during Pure PI (on trend), highlighting the relevance of this region in the absence of spatial cues and suggesting that adding a landmark reduces its involvement. This is in accordance with the idea that additional brain regions contribute to PI when spatial cues become available (21, 23).

In the right pmEC, we further demonstrated the existence of overall GLRs, but to control for potential order effects in our crossover design, we focused on differences between treatments on day one only. There, we found GLRs under placebo, but not under cortisol treatment, while control analyses specifically supported the presence of GLRs on day one only under placebo. This finding is in accordance with the hypothesis that cortisol affects the EC and impairs GLRs, thereby potentially inducing the several behavioral PI deficits that we found in a series of studies including the current one (31, 32). More specifically, because cortisol administration impairs inhibitory transmission in layer II of the EC (44), we propose that cortisol-induced inhibition of grid cells in the medial EC, an effect that correlates with the spacing of grid cells, causes behavioral PI deficits. Grid cells, and the medial EC, are generally related to PI performance, and lesions in the medial EC impair PI in rats (45, 46), presumably related to impaired distance estimations (47). Our analyses showed a relationship between GLRs and PI performance that depended on path distance, suggesting a beneficial role of GLRs for short incoming distances, but a detrimental role of GLRs over long incoming distances conditions. Based on previous studies (48, 22, 17), we assumed that GLRs should facilitate performance specifically in environments with no spatial cues, while our findings suggest that path distance is the more relevant moderator of GLR effects. In line with our finding, several lines of research indicate that grid cells might support PI specifically in case of short distances, including the emergence of grid distortions in larger environments (49), and the proposition that grid cells might rather provide a narrow and local rather than a general and global spatial metric (50). Over long incoming distances, grid distortions might lead to increased error accumulation, making errors larger and more likely to occur, a common observation in PI tasks, especially when no recalibration or anchoring of grid cells is possible. However, while this explanation would imply that grid cells are non-relevant over long distances, it does not cover the observation that they are detrimental, necessitating further research on this aspect.

A few limitations of this study need to be addressed. First, due to increased power, we used a within-subject crossover design. This can lead to results, where treatment and order effects cannot be disentangled, which we accounted for at the price of reducing sample size for GLR analyses. Also, stress effects in within-subject tasks differ from those in between-subject designs (51), even though we bypassed this issue by considering day one only (making it a pseudo between-subject design) for the stress effect on GLRs. Second, cognitive fatigue can water down stress effects on cognition (36, 51), and thus, the cognitive battery implemented before the task and the long duration of the task might have influenced our results, suggesting that cortisol effects on GLRs and PI could be even stronger than what we report here. Last, our sample consisted of young healthy men, which is not representative for the entire population, because gonadal hormones play a major in moderating effects of stress and cortisol (52). However, we have shown cortisol effects on PI in an older, female sample (32), and thus we can infer that behavioral effects of stress might be similar between sexes, while the investigation of neuronal effects is warranted in more representative samples in future studies.

### 3.1 Conclusion

Our current findings provide essential information about the underlying mechanisms of stress effects on PI, supporting the hypothesis that they are strongly mediated by effects of cortisol on the EC. Cortisol impaired overall PI performance in healthy human participants, while navigational patterns were altered towards navigating further away from the landmark. The fMRI data further demonstrated that cortisol increases activation in right CN in the presence of landmarks and impairs GLRs in the right pmEC. The GLRs correlated with better PI performance particularly over short distances.

## Acknowledgements

We thank Lukas Kunz for developing the experimental task and providing guidance for data acquisition. We also thank Liv Hog, Miriam Seyfried, Leja Kornefell and Leander Fester for their help with participant recruitment and data acquisition. Moreover, we thank Tobias Otto and PHILIPS Germany for technical support.

## Supplemental Information

Document S1. Figures S1-S5 and Tables S1-S5.

## Materials and Methods

### EXPERIMENTAL MODEL AND STUDY PARTICIPANT DETAILS

#### Participants

We used a two-day within-subject crossover design and recruited 42 healthy men, two of which had to be excluded due to technical failures of the joystick (see 2.5) and another one due to unsuccessful experimental manipulation (no cortisol increase, see 2.5), leaving a final sample size of *n* = 39, aged 19 – 34 years (24.18 ± 4.55 years; mean ± SD) and with a body mass index ranging between 18 and 29 kg/m² (23.95 ± 2.54 kg/m²; mean ± SD). We recruited participants through online advertisements in social media networks, mailing lists and in university classes at Ruhr University Bochum. Prior to testing, participants were instructed about all study procedures and gave written informed consent. The study was conducted in accordance with the Declaration of Helsinki as approved by the medical ethics committee of Ruhr University Bochum.

### METHOD DETAILS

#### Experimental design

Sample size was based on previous studies investigating PI (17), the functional relevance of GLRs (21, 22), and effects of cortisol administration (e.g., 53). Exclusion criteria comprised an acute or history of disease (i.e., neurological, psychiatric, cardiovascular, immunologic), current or history of medical or psychological treatment, drug use, female sex, previous experience with the PI paradigm (see experimental task), or any contraindication for participation in MRI studies. We only included men because of the well-known influence of sex hormones on the secretion of stress hormones (54) and the related changes in neural functioning (55). All participants had normal or corrected-to-normal vision and received a compensation of 10€/hour (60-70€ in total) or course credits.

Participants received two tablets of 10 mg cortisol (hydrocortisone, Hoechst) on one testing day, and two visually identical placebos on the other testing day in a double-blind two-day within-subject crossover design. Order of conditions was counterbalanced across participants. The dosage of 20 mg was chosen in accordance with previous studies demonstrating effects of cortisol on behavioral and neuronal responses with similar dosages (56–60). We assessed salivary cortisol using Salivettes (Sarstedt, Nümbrecht, Germany) collected at several time-points (see Fig. 2). Saliva samples were stored at -20 °C until assayed. Salivary cortisol concentrations were extracted from the samples using a time-resolved fluorescence immunoassay (IBL, Hamburg, Germany) at the local biochemical laboratory of Ruhr University Bochum and are reported in nanomole per liter (nmol/l). Intra- and inter-assay coefficients of variations were below 6.1%.

#### Experimental task

The Apple Game is a virtual PI task that was implemented by Bierbrauer et al. (22) via Unreal Engine 4 (Epic Games, version 4.11; see also Fig. 1 & Fig. S1). In this task, PI is considered as the ability to integrate across several paths and to calculate home-coming vectors based on visual cues only. A circular arena formed by an endless grassy plain with a blue sky rendered at infinity, with a diameter of 13,576 virtual meters (vm), built the environment. Briefly, each trial was composed of three phases. In the “start phase”, participants first moved to a basket and memorized its location (goal location). Then, the “outgoing phase” followed, in which they navigated to a variable number of trees (1–5), which appeared consecutively in different locations, until a tree containing a red apple (retrieval location) was reached. The variable number of trees allowed manipulating the path distance of the outgoing phase and thus difficulty. Basket and trees disappeared upon arrival at the respective locations. During the “incoming phase”, participants lastly were asked to take the shortest path from the retrieval location back to the goal location. They pressed a button when arriving at the presumed location (response location) and received visual feedback via zero to three stars corresponding to the Euclidean distance between response location and goal location (drop error; three stars for < 1,600 vm, two stars for < 3,200 vm, one star for < 6,400 vm). To investigate the influence of spatial cues on PI, the original version of the task contained three subtasks, of which we used two in this study. The “Pure PI” subtask was composed only of a grassy plain, forcing participants to solely rely on visual flow, while the “landmark-supported PI” (Landmark PI) subtask contained a central lighthouse as spatial cue. Participants first played up to eight training trials to get familiar with the task, during which a structural MRI scan (∼ 6 min) was obtained (see 2.6). Then, a total of 64 experimental trials (32 per subtask) divided into four blocks of 16 trials were presented, during which functional MRI scans were assessed (see 2.6). The outgoing phase of the 32 trials of each subtask was composed of six trials with 1, 2, 4 and 5 trees, respectively, and eight trials with three trees in randomized order. Before every new trial, a fixation cross was presented with a variable duration 5 – 7.5 s (randomly distributed).

#### Experimental Procedure

Testing sessions were performed after 1 p.m. to avoid interference with the cortisol awakening response (61). On day one, upon arrival, participants read study information and gave written informed consent, before undergoing a cognitive test battery, after which the pharmacological intervention and a series of questionnaires followed (see Tab. S5). About 30 minutes after pharmacological intervention, participants were prepared for the fMRI session, during which the experimental PI task was conducted. Lastly, participants answered a final set of questionnaires. The testing session on day two followed one week later, and the procedure was essentially the same, except for the absence of a cognitive test battery and the respective other pharmacological intervention. Finally, participants were debriefed and compensated (see Fig. 2).

#### Behavioral data

To extract Apple Game data from computer-generated log-files we used MATLAB (2021b, The MathWorks Inc., Massachusetts, US), including the Parallel Computing Toolbox (v6.12) and the CircStat Toolbox (62). Statistical analyses were conducted in R 1. (63) using the lme4 (64), lmerTest (65) and emmeans (66) packages.

#### MRI data

We acquired MRI data during the whole experimental task at the Bergmannsheil hospital in Bochum using a 3T Philips Achieva Scanner (Best, The Netherlands) with a 32-channel head coil. High-resolution whole-brain structural brain scans were acquired using a T1-weighted sequence with a 1 mm isotropic resolution, an FOV of 240 mm x 240 mm, and 220 transversally oriented slices during a total acquisition time (TA) of six minutes and three seconds. Blood oxygenation level–dependent (BOLD) contrast images were registered with a T2*-weighted gradient echoplanar imaging sequence with 2.5 mm isotropic resolution, TR = 2500 ms, TE = 30 ms, FA = 85°, FOV = 96 mm x 96 mm, 45 transversal slices in ascending order without slice gap, and TA = 17.32 ± 2.01 mins (mean ± SD), corresponding to 415.67 ± 48.27 (mean ± SD) volumes. Variation in TA emerged because the task was self-paced, leading to different durations of runs. In addition to three dummy scans preceding data acquisition, the first five images of each session were discarded to allow for signal steady-state transition. We further used a shim box of 60 mm x 60 mm x 60 mm around the medial temporal lobe to increase temporal signal-to-noise ratio. The virtual environment was presented to participants via MR-compatible liquid crystal display goggles (VisuaStim Digital, Resonance Technology Inc., Northridge, CA, USA) with a resolution of 800 × 600 pixels, and they navigated within the virtual environment using an MR-compatible joystick (Nata Technologies, Coquitlam, Canada).

fMRI data were preprocessed and subsequently analyzed using SPM12 implemented in MATLAB (2021a, The MathWorks Inc., Massachusetts, US) and nilearn for Python (Nilearn contributors, 2025). Preprocessing included slice time correction and spatial realignment. For whole-brain and ROI analysis, fMRI scans were further normalized to MNI space using parameters from the normalization procedure of the segmented structural T1 image. Further, we applied spatial smoothing with a 5 mm isotropic Gaussian kernel to the normalized fMRI data.

We used masks of the left and right HC, left and right CN, and bilateral PC (maximum probability masks; probability threshold: 0.25; Harvard-Oxford Cortical and Subcortical Structural Atlases, Harvard Center for Morphometric Analysis; https://cma.mgh.harvard.edu/). Because the medial EC is especially related to grid-cell activity in rodents and the human EC consists of structurally and functionally distinct subparts, where the anterior-lateral EC (alEC) and the posterior-medial EC (pmEC) represent the homolog of rodent lateral and medial EC, respectively, we focused our analysis on left and right pmEC using masks created by Maass et al. (67).

### QUANTIFICATION AND STATISTICAL ANALYSIS

#### Behavioral data analysis

One participant did not show any cortisol increase after cortisol administration and thus was excluded from statistical analyses, leading to a sample size of *n* = 39 for behavioral analyses. We assessed several measures for path distance, PI performance and navigational pattern during the experimental task. For path distance, two parametrical measures were assessed: O*utgoing distance* represents the accumulated distance from the goal to the retrieval location, while *incoming distance* refers to the Euclidean distance between retrieval and goal location (Fig. 1C). These measures represent different subcomponents of PI: Outgoing distance is relevant for keeping track of the travelled path in relation to the starting point (i.e., the later goal location), and incoming distance for calculating a direct vector to this goal location. Because previous studies indicated incoming distance to be more closely related to EC activation (68, 42), and the results of our previous work supported this view (32, 31), we again concentrated on this measure as a proxy of path distance. Moreover, overall PI performance is represented by the drop error, i.e., the Euclidean distance between response location and goal location (Fig. 1D). Lastly, to investigate the specific role of the landmark under heightened cortisol levels, we assessed two variables only applicable in Landmark PI. The first of these reflects the distance between the goal and the spatial cue (goal-to-landmark distance), while the other represents the mean Euclidean distance of the moving participant from the landmark across all time points of the incoming phase (movement-to-landmark distance).

#### fMRI data analysis

We excluded three participants, whose ECs were only partly covered, most likely due to susceptibility artifacts and other distortions, which are very strong in the medial temporal lobe (69), leaving a final sample size of *n* = 36 for the fMRI data analysis. In the first GLM, we modeled start phase, outgoing phase, incoming phase, and feedback separately for each of the two subtasks and each of the two treatments (16 regressors in total). The duration of these regressors corresponded to the duration of the respective phases in each trial. We tested three contrasts (plus their respective reverse contrasts): “Landmark PI > Pure PI”, “CORT > PLA” and their interaction “(Landmark PI > Pure PI)_CORT_ > (Landmark PI > Pure PI)_PLA_”, specifically during the outgoing and incoming phases, which represent the actual PI task. All regressors were convolved with the hemodynamic response function before entering the GLM. As nuisance regressors, we included motion parameters as estimated in the realignment procedure and a high-pass filter (time constant = 128 s) was implemented. We performed whole-brain analyses on each contrast. Contrast images from the first-level analysis of each participant were entered into a GLM. Statistical parametric maps were initially thresholded at a family-wise error (FWE)–corrected α level of *p* < 0.05 across the whole brain. We considered clusters significant at *p* < 0.05, FWE-corrected (extent threshold of five voxels). For all significant clusters, we provide maximum probability tissue labels with MNI coordinates derived from the Neuromorphometrics atlas as implemented in SPM12 (www.oasis-brains.org/; http://neuromorphometrics.com/). For ROI analyses, the significance threshold was set to *p* < 0.05 on voxel level, corrected for multiple testing within each ROI (FWE-corrected; using the small volume correction option of SPM12).

In a second GLM, we conducted an exploratory parametric modulation analysis of the drop error, which included one single unmodulated trial regressor as well as one parametric modulation regressor for trial-wise drop error modeling the combined outgoing and incoming phases of each trial. The onsets of these regressors corresponded to the onset of the outgoing phase and the modeled durations corresponded to the added total durations of the outgoing and incoming phases of each trial. We did not include separate regressors for each subtask (Landmark PI/Pure PI) since we were interested in the modulation of neural activity by PI performance across subtasks in this analysis. Drop error values were symmetrically inverted so that higher values of the parametric regressor would reflect better task performance. In addition, drop error values were normalized to range between 0 and 1 and mean-centered. Both regressors were convolved with the hemodynamic response function before entering the model. We included motion parameters resulting from realignment of functional images during preprocessing as nuisance regressors. Additionally, we used a cosine model with a high-pass frequency of 0.01 Hz to account for slow temporal drifts in the signal. Beta maps for the contrast of drop error parametric modulation against baseline were computed for each participant and each session. We performed a whole-brain analysis on the first-level drop error parametric modulation contrast images. Contrast images from each participant and each session were entered into a GLM which modeled the average of the first-level parametric modulation contrast across participants while accounting for subject-specific effects. Here, statistical parametric maps were thresholded at a false discovery rate (FDR)-corrected α level of *p* < 0.01 across the whole brain. Clusters were considered significant at *p* < 0.01, FDR-corrected and extending a minimum cluster size of ten voxels. For all significant clusters, maximum probability tissue labels are reported based on the peak voxel’s MNI coordinates and segmentations derived from the Harvard-Oxford atlas as implemented in nilearn. For the ROI analysis of drop error parametric modulation, average *p* values were corrected for the number of repeated tests using FDR-correction at α < 0.01.

To assess GLRs during fMRI, we performed a representational similarity analysis (RSA), using an approach analogous to previous studies (33, 22). This method assumes that grid cells in EC, due to their sixfold rotational symmetry, show similar activity patterns during movements that differ by 60° in angular space. Activity patterns should thus show higher similarity for movements with an offset of *n*\*60°, where *n* = {0, 1, …, 6}, and lower similarity for movements with an offset of *n*\*60° + 30°, where *n* = {0, 1, …, 5}.We refer to the first condition as “mod(α,60°) = 0°” and to the second condition as “mod(α,60°) = 30°” (where α represents the angular difference between movement directions, and “mod” denotes the modulo operator; see Fig. 5A). The RSA involved the following steps: (i) extracting the BOLD signal within the right pmEC ROI from the preprocessed fMRI volumes without normalization or smoothing; (ii) calculating the mean orientation and speed for each fMRI volume based on the trajectory data; (iii) excluding volumes where movement speed was slower than the first tertile; (iv) averaging volumes into bins of 5° orientation based on the mean orientation; (v) calculating the angular difference of the movement orientation between the volume bins; and (vi) calculating Fisher z-transformed Pearson correlations between the volume bins. We then compared the pattern similarity of pairs of volume bins offset by ±15° from the mod(α,60°) = 0° condition (aligned movements) with those offset by ±15° from the mod(α,60°) = 30 condition (misaligned movements). Pattern similarity was expected to be higher for aligned movements than for misaligned movements, thus reflecting GLRs. Two control analyses were performed. First, to ensure that the increase in pattern similarity was specific to sixfold symmetry, the same analysis was performed for other types of rotational symmetry (4-fold, 5-fold, 7-fold and 8-fold). Second, the analysis was also performed after excluding correlations that were ±15° from 0°, to ensure that increases in pattern similarity were not primarily driven by movements in the same direction, which could reflect head direction signals.

#### Statistical analysis

Our first statistical analysis examined the success of pharmacological intervention by comparing cortisol concentrations using a rANOVA with time and pharmacological intervention as within-subject factors. Because cortisol concentrations typically exhibit a right-skewed distribution, we conducted a natural log (ln) transformation to obtain normally distributed data. In case of violation of the sphericity assumption, we used Greenhouse-Geisser adjustment (and for the sake of presentation, we rounded the corrected degrees of freedom to the nearest whole number). Post-hoc pairwise comparisons were performed using Bonferroni-corrected *t*-tests (or Welch’s *t*-tests in case of unequal variances).

To test the hypothesis postulating differential effects of cortisol depending on subtask and path distance on PI performance, we then built a linear mixed model with PI performance (i.e., drop error) on the level of single trials as criterion, and subtask (two levels: Pure PI vs. Landmark PI), path distance (i.e., incoming distance), and treatment (two levels: cortisol vs. placebo) as within-subject predictors (Model 1; Tab. S1).

To test the hypothesis concerning the effects of cortisol on the role of the landmark, we built two further linear mixed models (Models 2A-B; Tab. S1). The first of these aimed at investigating the role of goal-to-landmark distance and its interaction with cortisol on PI performance and thus included PI performance on the level of single trials as criterion, goal-to-landmark distance (only in Landmark PI), and treatment as within-subject predictors. The second of these models aimed at investigating whether cortisol affected the employment of navigational strategies (irrespective of performance) and included movement-to-landmark distance (only in Landmark PI) on the level of single trials as criterion, and treatment as within-subject predictor.

To evaluate the overall existence of GLRs, we conducted a one-tailed one-sample t-test comparing activation of EC during aligned vs. misaligned movements against zero (Model 3A; Tab. S1). To test whether GLRs were related to treatment, we conducted a linear model with GLRs as criterion, treatment and day as within-subject predictors, and age as covariate (Model 3B; Tab. S1). We conducted separate t-tests for the control analyses (other symmetries or correction for heading direction). In the next step, to examine the behavioral relevance of GLRs, we built a linear model with PI performance on the level of single trials as criterion, and subtask, path distance, and GLRs as within-subject predictors (Model 4; Tab. S1).

In all linear mixed models, “subject” was added as random factor, and age, testing day, and sequence (to control for order effects) as covariates. For all analyses, we centered within-subject parametric predictors (incoming distance, goal-to-landmark distance) to the participant’s mean and age to the grand mean of all participants (70). For analysis of fixed effects, we always used type III sum of squares. In case of interactions between parametric predictors, we discretized all but one parametric predictor and calculated estimated marginal means (“emmeans” or “adjusted means”) or estimated marginal means of linear trends (“emtrends” or “conditional regression equations”) based on the minimum, mean and maximum values of these discretized predictors. For post-hoc tests against zero or pairwise comparisons using Fisher’s tests, we used Šídák-adjustment correcting for number of discretized predictors (3), number of subtasks (2), number of treatments (2), or a combination of those. For an approximation of degrees of freedom, we used the Kenward-Roger method (and for the sake of presentation, we rounded them to the nearest whole number). To estimate effect sizes, we used Partial Eta-Squared (η_p_^2^) for *F*-tests and Cohen’s *d* for *t*-tests. Multicollinearity between predictors was not problematic (all variance inflation factors < 5). Assumptions of statistical tests were verified before application and all tests were conducted two-tailed (except for t-tests on GLRs) at a significance level of α = 0.05.

