## Supplemental Material for "The relationship between cortisol, grid-like representations and path integration"

### Supplementary Material

#### Supplementary Figures

##### A Bird‘s eye view of virtual environment

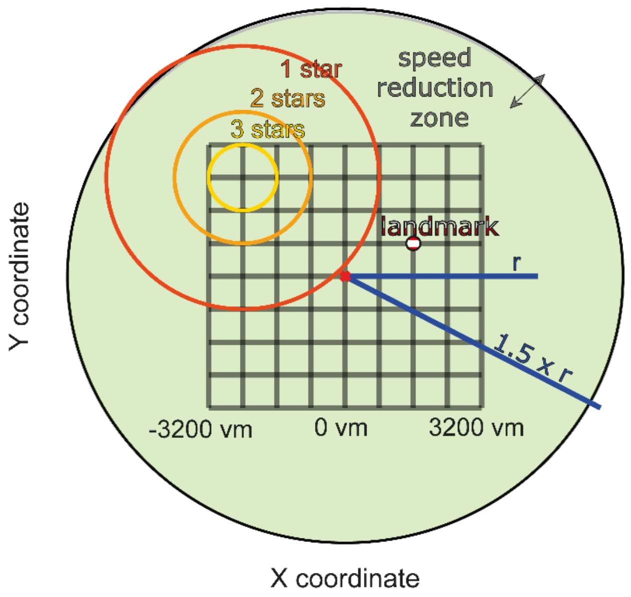


##### B Examples for the assignment of locations to the grid

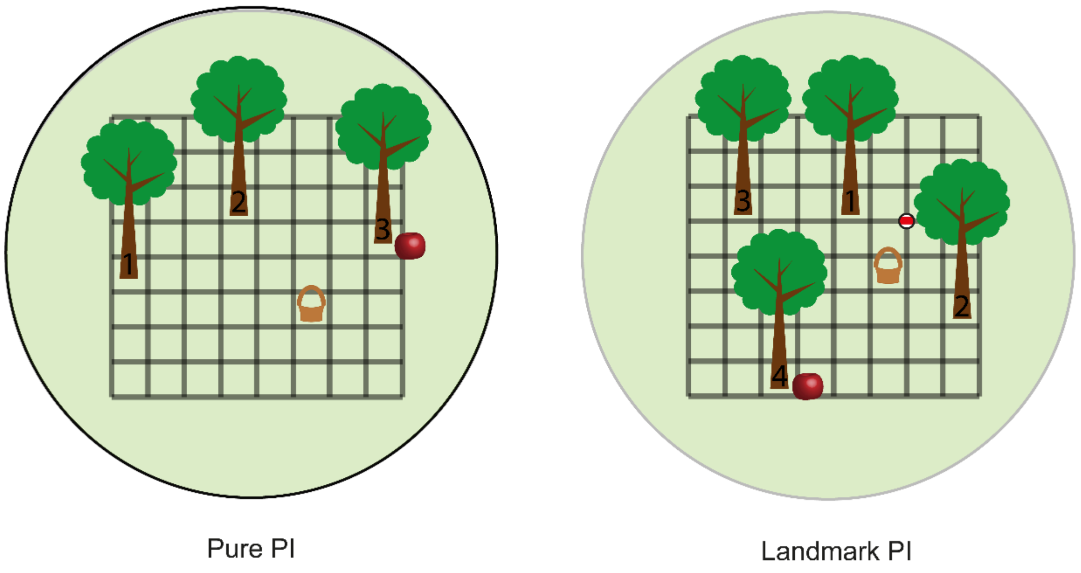


**Figure S1. Overview of the virtual environment. (A)** Locations of baskets (i.e., goal locations) and trees were equally distributed across a grid of 8x8 squares such that all participants visited all squares at least once in each subtask. Feedback was given according to the Euclidean distance between the response location and the correct goal location (i.e., drop error). In all subtasks, participants’ speed was linearly decreased to zero when their distance from the center of the arena was larger than 1.25*r vm. In Landmark PI, a landmark was located close to the center of the environment (at x = 1600 vm, y = 800 vm). **(B)** Two examples of how locations could be assigned to the grid. Numbers represent the order at which the specific trees appeared. Pure PI: pure path integration, Landmark PI: landmark-supported path integration, r: radius, vm: virtual meters. Figure adapted from Bierbrauer et al. (2020).

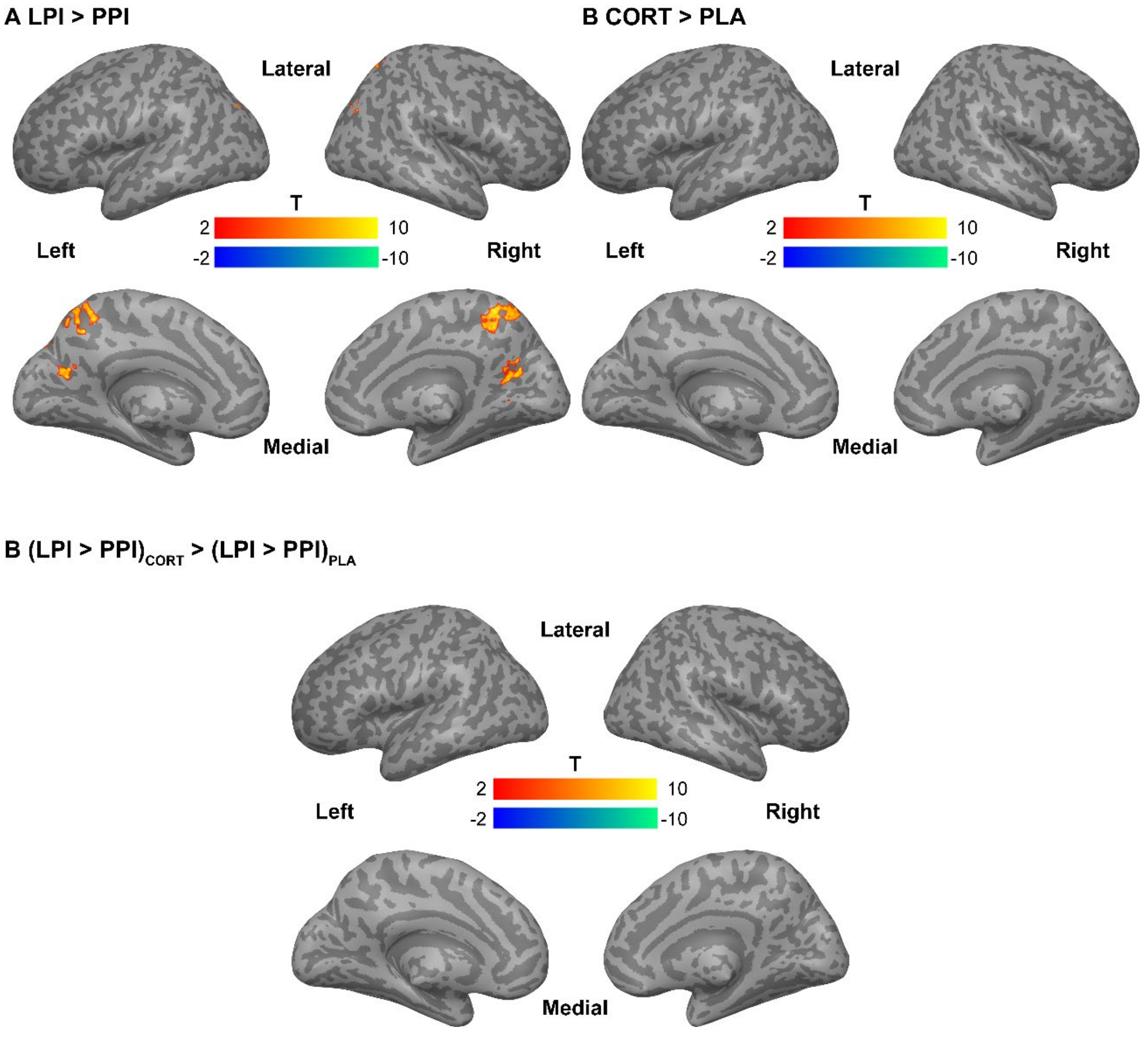


**Figure S2. Whole-brain analysis***.* Depicted are clusters with more than 5 voxels surviving an initial height threshold of $p < .05$, FWE-corrected for whole brain. For significant clusters, maximum probability tissue labels were derived from the Neuromorphometrics atlas in SPM. (A) Clusters in R precuneus, L precuneus, R middle occipital gyrus, L superior occipital gyrus and L superior parietal lobule survived statistical correction. (B) No cluster survived statistical correction. (C) No cluster survived statistical correction LPI: landmark-supported path integration, PPI: pure path integration.

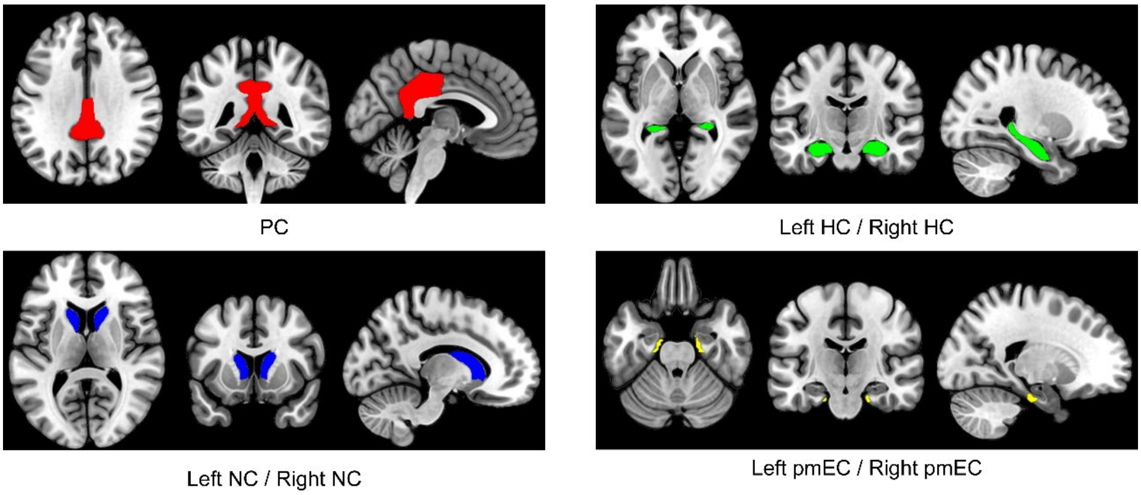


**Figure S3. Masks used for ROIs.** We used masks for the left and right pmEC (top-left panel), the left and right HC (top-right panel), the PC (bottom-left panel) and the left and right NC (bottom-right panel). PC: posterior cingulate, HC: hippocampus, NC: nucleus caudate, pmEC: posterior-medial entorhinal cortex.

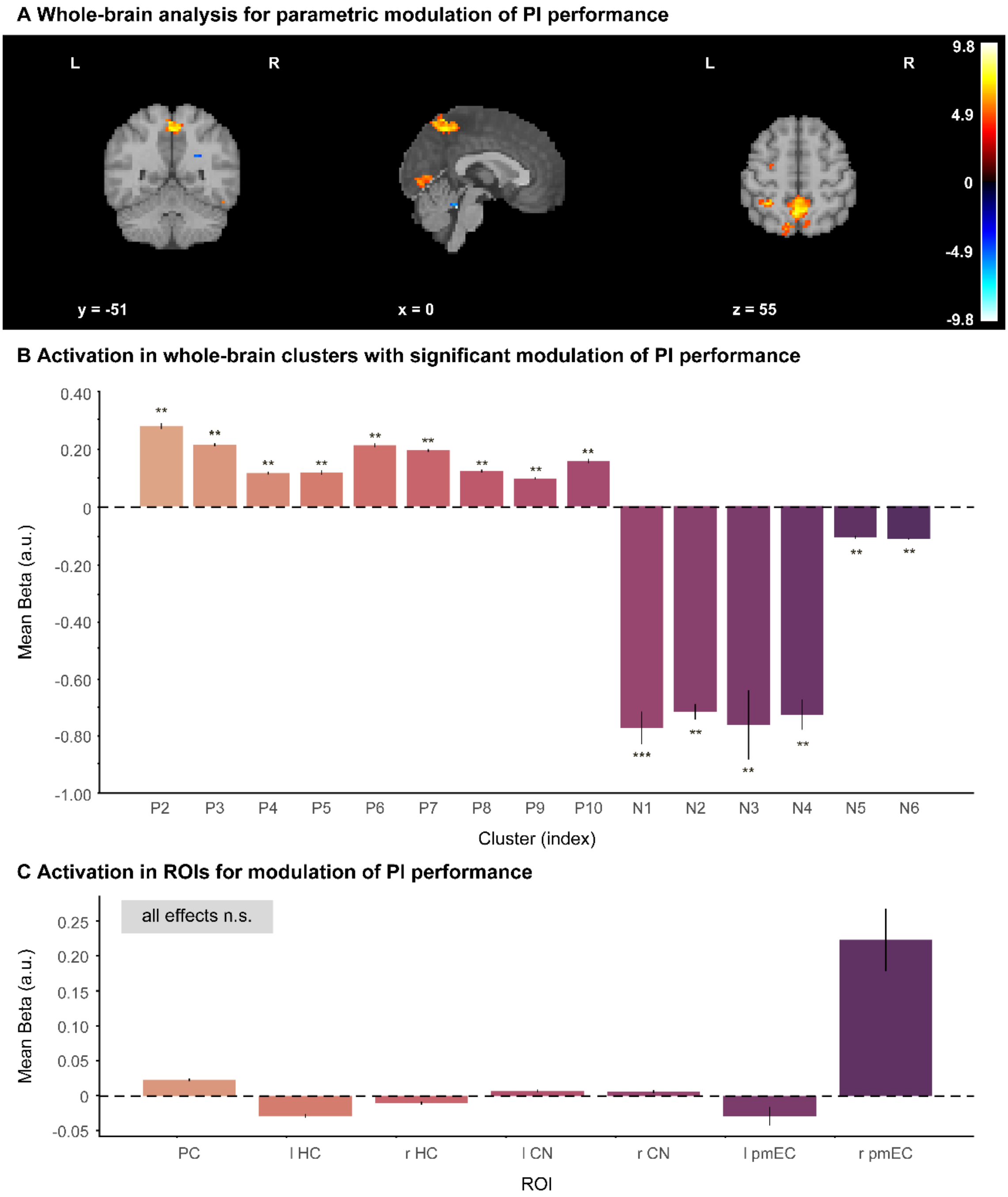


**Figure S4. Parametric modulation of PI Performance**. (A) Depicted are clusters with more than 10 voxels surviving an initial height threshold of p < .01, FDR-corrected for whole brain. For significant clusters, maximum probability tissue labels were derived from the Harvard-Oxford atlas in nilearn. Brain plots show color-coded t-values for the average contrast of drop error parametric modulation against baseline across participants. (B) Activation in whole-brain clusters with significant positive modulation (increased activation with higher performance in a given trial, P2 – P10) and negative modulation (reduced activation with higher performance in a given trial, N1 – N6) by inverted drop error (effects across subtasks and treatments). Cluster indices correspond to brain regions listed in Tab. S4. (C) FDR-correction was applied for the number of ROIs at α < .01. No ROI showed significant modulation of the BOLD signal by (inverted) drop error. ROIs were tested for significant parametric modulation across subtasks and treatments. Bars depict mean beta estimates for the drop error parametric regressor in each cluster/ROI. Error bars represent SEM. l: left, r: right, PC: posterior cingulate, HC: hippocampus, CN: caudate nucleus, pmEC: posterior-medial entorhinal cortex, n.s.: not significant, ****p* < .001, ** *p* < .01.

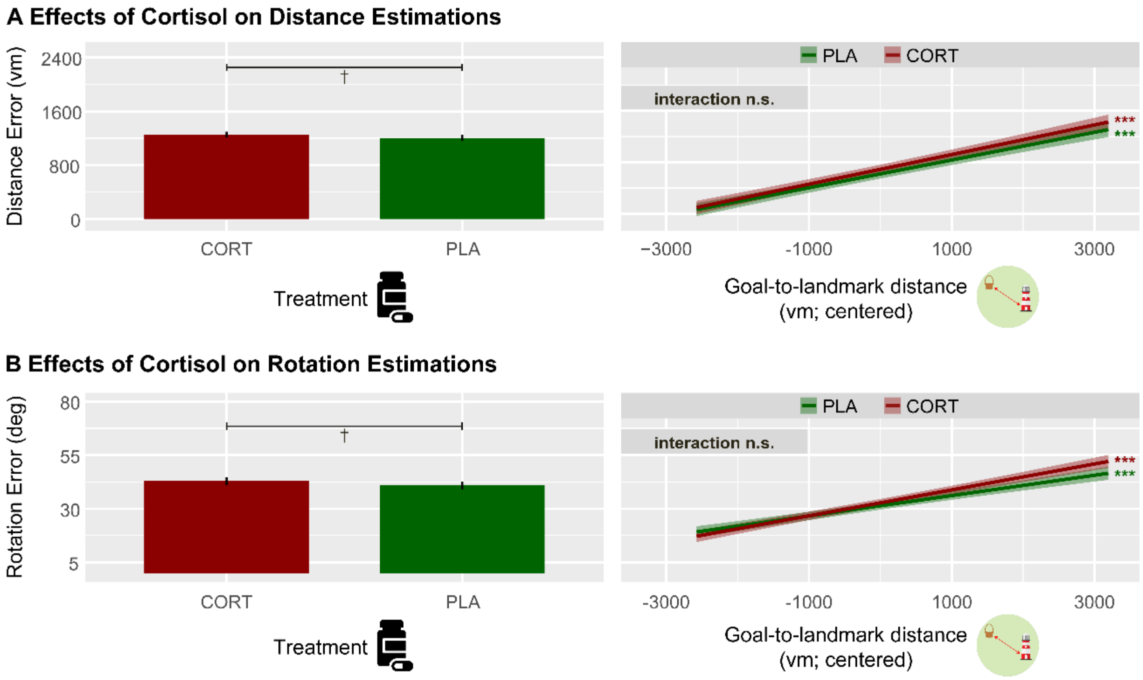


**Figure S5. Effects of cortisol on PI subcomponents**. (A) Cortisol (on trend) led to higher distance errors (left). In Landmark PI trials, a higher distance between goal location and landmark location predicted higher distance error and this effect did not differ between treatment conditions (right). (B) Cortisol (on trend) led to higher rotation errors (left). In Landmark PI trials, a higher distance between goal location and landmark location predicted higher rotation error and this effect did not differ between treatment conditions (right). Significant slopes are indicated via asterisks. Error bars and confidence bands represent SEM. CORT: cortisol, PLA: placebo, vm: virtual meters, n.s.: not significant, $^{***}p < .001$, $^{\dagger}\ p < .10$.

#### Supplementary Tables

Table S1. Statistical Models.

| Model | Criterion | Predictors (within Subject) | Predictors (between Subjects) | Covariates | Random Effect |
|---|---|---|---|---|---|
| **Effects of Subtask, Path Distance and Cortisol on PI Performance** | | | | | |
| **1** | Drop Error | Subtask, Incoming Distance, Treatment | - | Age, Day, Sequence | Subject |
| **Effects of Landmark (only LPI trials)** | | | | | |
| **2A** | Drop Error | Goal-to-Landmark Distance, Treatment | - | Age, Day, Sequence | Subject |
| **2B** | Movement-to-Landmark Distance | Treatment | - | Age, Day, Sequence | Subject |
| **Effects of cortisol on GLRs** | | | | | |
| **3A** | GLRs | - | - | - | - |
| **3B** | GLRs | Treatment, Day | - | Age | - |
| **Effects of Subtask, Path Distance, Cortisol and GLRs on PI Performance** | | | | | |
| **4** | Drop Error | Subtask, Incoming Distance | GLRs | Age | - |

*Note.* Overview of statistical models used to examine effects of cortisol administration on PI (Model 1), navigational pattern in the presence of a landmark (Models 2A–2B), GLRs (Models 3A–B), and to investigate the relationship between GLRs and PI (Model 4). PC: posterior cingulate, NC: nucleus caudate, pmEC: posterior-medial entorhinal cortex, PI: path integration, PPI: pure path integration, LPI: landmark-supported path integration, CORT: cortisol, PLA: placebo, GLRs: grid-like representations (in right pmEC).

Table S2. Differences in cortisol concentrations after pharmacological treatment.

| timepoint | placebo (n = 39) | cortisol (n = 39) |
|---|---|---|
| baseline | 3.55 ± 2.72 | 3.73 ± 2.60 |
| +30 mins | 2.80 ± 2.45 | 323.44 ± 338.44*** |
| +110 mins | 1.66 ± 1.05 | 57.54 ± 82.56*** |

*Note.* To verify the success of cortisol administration, we analyzed cortisol concentrations using a repeated-measures analysis of variance (rANOVA) with timepoint and pharmacological intervention as within-subject factors. We found significant main effects of treatment ($F_{(1,37)}$ = 246.15, $p$ < .001, $\eta_p^2$ = .869) and timepoint ($F_{(2,74)}$ = 171.22, $p$ < .001, $\eta_p^2$ = .822), and a significant timepoint x treatment interaction ($F_{(2,74)}$ = 205.07, $p$ < .001, $\eta_p^2$ = .847). Post-hoc pairwise comparisons revealed higher cortisol concentrations for both timepoints following cortisol compared to placebo administration (both $t \geq 16.28$, both $p_{Bonferroni} \leq .001$, both $d \geq 2.607$), i.e., for the entire PI task, whereas no differences occurred at baseline ($t_{(38)}$ = 1.01, $p_{Bonferroni}$ = 1, $d$ = 0.162). Cortisol concentrations represent mean ± standard deviation in nmol/l; $p$-values extracted from separate $t$-tests between treatments; ***$p$ < .001.

Table S3. Global and local maxima of whole brain analysis for contrasts.

| Contrast / Region | Voxels | X | Y | Z | *t*-score |
|---|---|---|---|---|---|
| Landmark PI > Pure PI | | | | | |
| R precuneus | 125 | 18 | -58 | 18 | 9.11*** |
| L precuneus | 541 | -2 | -52 | 52 | 8.03*** |
| L precuneus | 58 | -8 | -62 | 18 | 7.65*** |
| R middle occipital gyrus | 20 | 38 | -78 | 35 | 6.85*** |
| L superior occipital gyrus | 33 | -25 | -80 | 32 | 6.73** |
| L superior parietal lobule | 5 | -20 | 78 | 45 | 6.33* |
| **Posterior cingulate (SVC)** | **233** | **8** | **-50** | **5** | **6.65*** ** |
| **R nucleus caudate (SVC)** | **29** | **15** | **2** | **20** | **4.45*** |
| CORT > PLA | No significant clusters | | | | |
| (Landmark PI > Pure PI)$_{CORT}$ > (Landmark PI > Pure PI)$_{PLA}$ | | | | | |
| **R pmEC (SVC)** | **4** | **22** | **-20** | **-28** | **2.78*** |
| **R nucleus caudate (SVC)** | **177** | **10** | **8** | **12** | **4.62** ** |
| **L nucleus caudate (SVC)** | **156** | **-12** | **-5** | **20** | **3.90** ** |
| Pure PI > Landmark PI | No significant clusters | | | | |
| PLA > CORT | No significant clusters | | | | |
| (Landmark PI > Pure PI)$_{PLA}$ > (Landmark PI > Pure PI)$_{CORT}$ | No significant clusters | | | | |

*Note*. Brain regions exhibiting BOLD activations for the individual contrasts. Reported are all clusters with more than 5 voxels, surviving an initial height threshold of $p < 0.05$, FEW-corrected for whole brain, as well as small volume corrected (SVC; FWE corrected $p < 0.05$) clusters for pmEC, HC, caudate nucleus, and PC/RSC. Clusters within ROIs are marked bold. For other significant clusters, maximum probability tissue labels are derived from the Neuromorphometrics atlas contained in SPM. L, left; R, right, ****p* < .001, ***p* < .01, **p* < .05.

Table S4. Global and local maxima of whole brain analysis for drop error parametric modulation.

| Cluster No. | Region | Voxels | X | Y | Z | *z*-score |
|---|---|---|---|---|---|---|
| **Clusters showing positive modulation by inverted drop error** | | | | | | |
| P1 | Unknown | 12 | 50 | -62 | -47 | 8.81** |
| P2 | R Temporal Occipital Fusiform Cortex | 40 | 42 | -47 | -20 | 6.81** |
| P3 | Precuneus Cortex | 303 | -25 | -57 | 57 | 6.50** |
| P4 | L Superior Parietal Lobule | 32 | -30 | -45 | 55 | 5.63** |
| P5 | L Supramarginal Gyrus, anterior division | 22 | -62 | -32 | 37 | 5.49** |
| P6 | L Lateral Occipital Cortex, superior division | 50 | -10 | 70 | 52 | 5.29** |
| P7 | Medial Lingual Gyrus | 83 | 25 | -77 | 0 | 5.16** |
| P8 | L Middle Frontal Gyrus | 69 | -35 | 37 | 32 | 5.08** |
| P9 | L Precentral Gyrus | 16 | -27 | -75 | 60 | 4.99** |
| P10 | R Supramarginal Gyrus, anterior division | 22 | 62 | -30 | 45 | 4.90** |
| **Clusters showing negative modulation by inverted drop error** | | | | | | |
| N1 | R Lateral Occipital Cortex, inferior division | 16 | 32 | -87 | -5 | -9.15*** |
| N2 | L Occipital Fusiform Gyrus | 39 | -30 | -82 | -12 | -7.44** |
| N3 | Unknown | 10 | 7 | -35 | -27 | -7.12** |
| N4 | Unknown | 33 | 0 | -40 | -25 | -6.64** |
| N5 | Unknown | 29 | 15 | -47 | 25 | -5.60** |
| N6 | Unknown | 35 | -35 | -40 | 0 | -5.18** |

*Note*. Brain regions exhibiting BOLD activations significantly modulated by trial-wise (inverted) drop error. Reported are all clusters with more than 10 voxels, surviving an initial height threshold of $p < 0.01$, FDR-corrected for whole brain. Maximum probability tissue labels are derived from the Harvard-Oxford atlas as implemented in nilearn. L, left; R, right, ***$p < .001$, **$p < .01$.

Table S5. List of cognitive tests and questionnaires.

| Cognitive test | Reference | Cognitive dimension |
|---|---|---|
| Ray-Figure Task | Rey (1941) | Visual memory |
| Logical memory | Wechsler (2008) | Verbal memory |
| Digit span test | Wechsler (2008) | Verbal working memory |
| Block span test | Corsi (1972) | Visual working memory |
| Trailmaking A & B | Reitan (1955) | Visual attention and Task switching |
| **Questionnaire** | **Reference** | **Measured variable** |
| Positive and Negative Affect Schedule | Krohne et al. (1996) | Positive and negative affect |
| Edinburgh Handedness Inventory | Oldfield (1971) | Laterality quotient |
| Navigation Strategy Questionnaire | Zhong and Kozhevnikov (2016) | Navigational strategy preferences |
| Life Threatening Experiences | Brugha and Cragg (1990) | Traumatic experiences in the last 12 months |
| Big Five Inventory K | Rammstedt and John (2005) | Big five personality dimensions |
| Depression-Anxiety-Stress Scale | Nilges and Essau (2015) | Depression, anxiety, and stress levels |
| Apple Game Questionnaire | - | Strategy use during Apple Game |

*Note.* Cognitive tests were only conducted on day 1. These results are not part of this study.
